# Parthenogenote-Derived Brain Unveils the Critical Role of Paternal Genome in Neural Development

**DOI:** 10.64898/2026.01.06.697935

**Authors:** Marina Takechi, Yezhang Zhu, Zezhen Lu, Ying Zeng, Ken-Ichi Mizutani, Toru Nakano, Li Shen, Shinpei Yamaguchi

## Abstract

Genomic imprinting, an epigenetic mechanism that governs parent-of-origin–specific gene expression, is essential for mammalian development, yet its role in late-stage development remains unclear due to the lethality of parthenogenetic (Pg) embryos. Here, we establish cell replacement with parthenogenote-derived cells (CReP), a blastocyst complementation strategy that enables survival and tissue-specific contribution of Pg-derived cells. Brain-targeted CReP showed that Pg-derived cells can participate in neural development but fail to maintain neuronal–glial balance due to aberrant activation of Notch signaling caused by the loss of the paternally expressed gene Dlk1. Restoration of Dlk1 normalized Notch activity and rescued neuronal differentiation. These findings reveal a critical role of the paternal genome, through Dlk1-mediated regulation of Notch signaling, in ensuring neural stem cell expansion and balanced cell fate decisions. The CReP model provides a powerful platform for investigating genomic imprinting and parental genome contributions in development and disease.

## Introduction

Genomic imprinting is an epigenetic mechanism that governs parent-of-origin-specific gene expression. First proposed in the 1980s, the concept emerged from pronuclear transplantation studies, which revealed uniparental embryos failed to develop normally[1]. These findings provided the compelling evidence that maternal and paternal chromosomes have distinct functional roles in mammalian development. Whereas parthenogenesis̶the development of offspring from an unfertilized egg̶is widespread in diverse species, including invertebrates, fish, amphibians, birds, and reptiles[2]. In mice, parthenogenetic (Pg) embryos, containing only maternal genomes, exhibit severe placental development defects and fail to survive beyond embryonic day 9.5 (E9.5). Androgenetic (Ag) embryos, containing only paternal genomes, display even more severe phenotypes[1,3]. These observations highlight the mammalian-specific nature of genomic imprinting, which likely evolved to support unique features such as the placenta and large brains[4,5].

Analysis of knockout mice for classical imprinted genes, such as *Igf2* and *Igf2r*, led to the proposal of conflict hypothesis (or kinship theory) of genomic imprinting[6–8]. This theory posits that genomic imprinting arose from an evolutionary conflict between maternal and paternal genomes over resource allocation to offspring. For instance, *Igf2-*knockout mice exhibit growth restriction, while *Igf2r-*knockout display overgrowth, illustrating the opposing effects of paternal and maternal gene expression[6,7]. The paternal genome is thought to prioritize maximizing offspring growth and resource acquisition. In contrast, the maternal genome tends to favor a balanced resource distribution to ensure maternal survival and future reproductive opportunities, a phenomenon particularly evident in placental mammals[9–13].

Approximately 200 imprinted genes have been identified in humans and mice[10,14], many of which are highly expressed in the placenta and brain[14]. Disruptions of their monoallelic expression cause severe disorders, such as Prader-Willi, Angelman, Beckwith-Wiedemann, and Silver-Russell syndromes[14–19]. Advanced maternal age and assisted reproductive technologies (ART) have also been associated with imprinting disorders[20]. Despite their importance, the physiological roles and regulatory mechanisms of imprinting remain incompletely understood.

Genomic imprinting is established by epigenetic modifications, particularly DNA methylation at differentially methylated regions (DMRs), which distinguish the maternal and paternal alleles. Germline DMRs are established during gametogenesis, whereas somatic DMRs acquire DNA methylation after fertilization under the control of germline marks[21]. In addition, certain imprinted genes exhibit tissue-specific imprinting, such as maternal expression of *Igf2* in the brain[22] [23]. Non-canonical mechanisms, such as maternal allele-specific H3K27me3, further illustrate the complexity of imprinting regulation[24].

Collectively, these findings underscore both the complexity of imprinting regulation and the necessity of advancing current methodologies to fully elucidate its scope.

In this study, we established a novel blastocyst complementation system to generate chimeric mice with brains composed of Pg-derived cells. This approach enables investigation of the distinct contributions of maternal and paternal genomes during later developmental stages that were previously inaccessible due to embryonic lethality. Our results demonstrate that the paternal genome is critical for neural cell fate determination, acting through Dlk1-mediated regulation of Notch signaling. This experimental framework provides a powerful platform for exploring the indispensable roles of parental alleles in embryogenesis and offers new insights into the mechanisms of genomic imprinting in mammals.

## Results

### Establishing CReF and CReP via *Wnt1*-KO blastocyst complementation

A straightforward way to interrogating the physiological roles of genomic imprinting is to analyze of uniparental embryos, which inherit alleles from a single parent. However, the early lethality of Pg and Ag embryos shortly after implantation restricts their use at later developmental stages[1,3]. To overcome this limitation and investigate allele-specific functions at later stages, we first generated chimeric embryos by aggregating Pg and fertilized embryos (Supplementary Fig. S1a). At E9.5, chimeric embryos with high contribution of Pg-derived cells exhibited reduced body size and developmentally delayed relative to controls (Supplementary Fig. S1b). By E14.5, Pg cell contribution was minimal, suggesting the progressive elimination of Pg-derived cells̶likely via cell competition or embryonic lethality in chimeras with substantial Pg contributions (Supplementary Fig. S1c).

To enhance Pg-derived cell contribution at later stages, we employed a blastocyst complementation method[25], using gene-knockout (KO) fertilized embryos as recipients. Specifically, we targeted the brain by knocking out *Wnt1*, a gene essential for the midbrain and cerebellum development[26]. Genotyping confirmed mutations in the *Wnt1*-KO embryos, and histological analysis verified the expected brain agenesis phenotype (Supplementary Figs. S2A to S2C, Supplementary Table S1). Aggregation of wild-type with *Wnt1*-KO embryos demonstrated that GFP-positive wild-type cells efficiently complemented brain development in the chimeric embryos, contributing broadly across the midbrain and cerebellum (Supplementary Figs. S3a to S3d). Furthermore, chimeric neonates were born at term without gross morphological abnormalities and developed normally to adulthood (Supplementary Fig. S3e). We designated this method as Cell Replacement with Fertilized embryo-derived cells (CReF) and resulting mice as CReF mice.

Next, we generated chimeras by aggregating Pg embryos with *Wnt1*-KO recipients.

Compared with chimeras using fertilized recipients̶which showed only minimal Pg-derived cell contributions—*Wnt1*-KO chimeras exhibited significant GFP-positive Pg-derived cell contributions to the brain at E14.5 (Fig. 1a and 1b). We termed this approach Cell-Replacement with Parthenote-derived cells (CReP) and resulting mice as CReP mice.

**Fig. 1.**
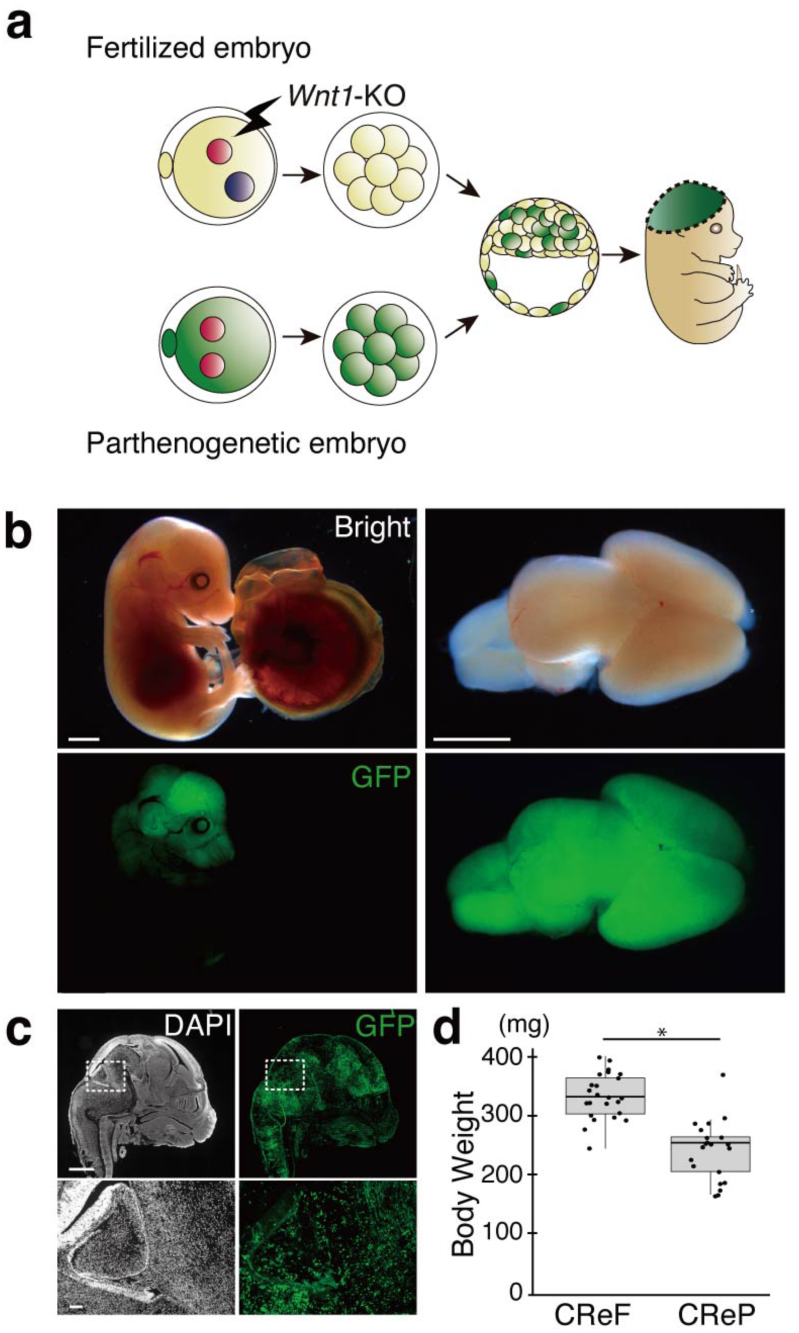
Parthenogenetic (Pg) donor complementation (CReP) enables late-stage development and rescues neural morphology while growth remains restricted. (**a**) Experimental scheme for establishing Cell-replacement with Parthenogenote derived cells (CReP) embryos. (**b**) Representative images of E14.5 CReP embryos. GFP-positive cells indicate Pg-derived contribution. Scale bars, 1mm. (**c**) Representative cryosection images of the E14.5 CReP head. scale bar, 1 mm. (**d**) Box plot showing body weight of E14.5 CReF and CReP embryos. Each dot indicates an individual measurement. CReF: n = 24. CReP: n = 16. \**P*<0.05

Remarkably, as with CReF embryos, the morphological defects observed in *Wnt1*-KO embryos were fully rescued in CReP embryos (Fig. 1c). However, at E14.5, CReP embryos exhibited significantly lower body weight than CReF counterparts (Fig. 1d), and most CReP neonates died shortly after birth. (Supplementary Figs. S4a to S4c).

### CReP identifies candidate imprinted genes in the embryonic brain at later developmental stages

CReP allowed Pg-derived cells to bypass early developmental barriers, providing a platform to investigate the consequences of paternal-allele absence at later developmental stages. We profiled gene expression in the embryonic brain by RNA-sequencing (RNA-seq) of GFP-positive donor cells purified by fluorescence-activated cell sorter (FACS). Because Pg-derived cells lack paternal alleles, we expected maternally expressed genes (MEGs) to show approximately twofold higher expression relative to wild-type controls, while paternally expressed genes (PEGs) would be essentially absent. To avoid missing candidates, we applied permissive thresholds in the initial screen and identified 319 up-regulated genes (log2Fold change (FC) > 0.8, FDR < 0.05), and 28 down-regulated genes (log2FC < -2, FDR < 0.05) (Fig. 2A). An up-regulation trend (log2FC > 0, FDR < 0.05) was observed in 8.5% (5/59) of MEGs, compared with 1.7% (330/19760) of non-imprinted genes (Non-IGs) and none (0/47) of PEGs. Conversely, a down-regulation trend (log2FC < 0, FDR < 0.05) was seen in 38% (18/47) of PEGs, 3.4% (2/59) of MEGs, and 0.75% (149/19760) of Non-IGs (Fig. 2b).

**Fig. 2.**
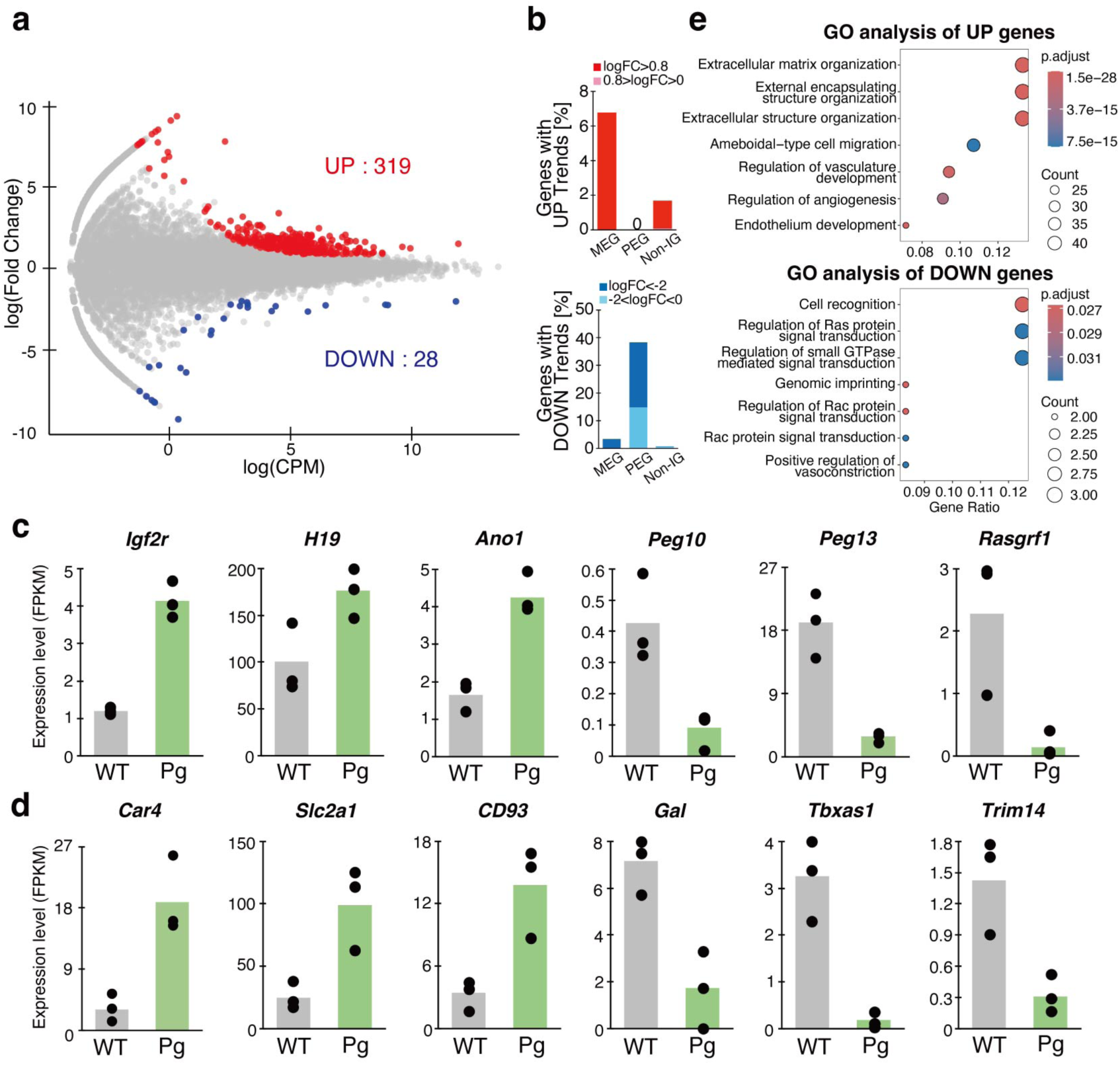
Gene expression profile of Pg-derived cells from CReP brains. (**a**) MA plot of GFP-positive sorted cells from E14.5 CReF and CReP brains. Up-regulated genes (red, log2FC > 0.8, FDR< 0.05); down-regulated genes (blue, log2FC < -2, FDR < 0.05). (**b**) Gene expression trends for known maternally expressed genes (MEGs), paternally expressed genes (PEGs), and non-imprinted genes (Non-IGs) among genes up-regulated or down-regulated in Pg-derived cells from CReP relative to WILD-TYPE donor cells from CReF brains. (**c and d**) Expression level of representative imprinted genes (**c**), and differentially expressed genes excluding known imprinted genes (**d**). (**e**) Gene ontology enrichment for up-regulated (Top), and down-regulated (Bottom) genes in Pg-derived cells.

Notably, among 83 known MEG cataloged in a previous study[27], four genes̶including *Igf2r*, *H19*, and *Ano1*—showed approximately two-fold higher expression in Pg cells from CReP than in wild-type cells from CReF embryos (Fig. 2c, Supplementary Table S2).

Conversely, eleven of the 66 known PEGs[27]̶including such as *Peg10*, *Peg13*, and *Rasgrf1* were significantly represses (Fig. 2c, Supplementary Table S2). We also identified 332 differentially expressed genes (DEGs) that may include previously uncharacterized imprinted gene candidates (e.g., *Car4, Slc2a1, CD93, Gal, Tbxas1,* and *Trim14*) (Fig. 2d, Supplementary Table S2). Gene ontology enrichment showed that up-regulated genes were enriched for extracellular matrix organization and external encapsulating structure organization, whereas down-regulated genes were enriched for cell recognition and regulation of Ras protein signal transduction (Fig. 2e, Supplementary Table S3). Collectively, these data illustrate that CReP enables systematic identification of imprinted genes.

### Widespread depletion of Pg-derived cells with reduced neural stem/progenitor populations in CReP brains

To investigate cellular heterogeneity and developmental abnormalities in CReP brains, we performed single-nucleus RNA sequencing (snRNA-seq) on brain cells from E14.5 CReP and CReF embryos. Clustering resolved 26 cell states, and in every cluster the fraction of GFP-positive donor-derived cells was lower in CReP brains than in CReF (Figs. 3a to 3c, Supplementary Table S4). This pervasive reduction indicates a generalized competitive disadvantage of Pg-derived cells across neural lineages during midgestational brain development. Notably, the proportion of Pg-derived GFP-positive cells was lowest in Cluster 1, representing a neural stem cell population characterized by the expression of markers such as *Meg3, Snhg11* and *Csmd3* (Fig. 3b).

**Fig. 3.**
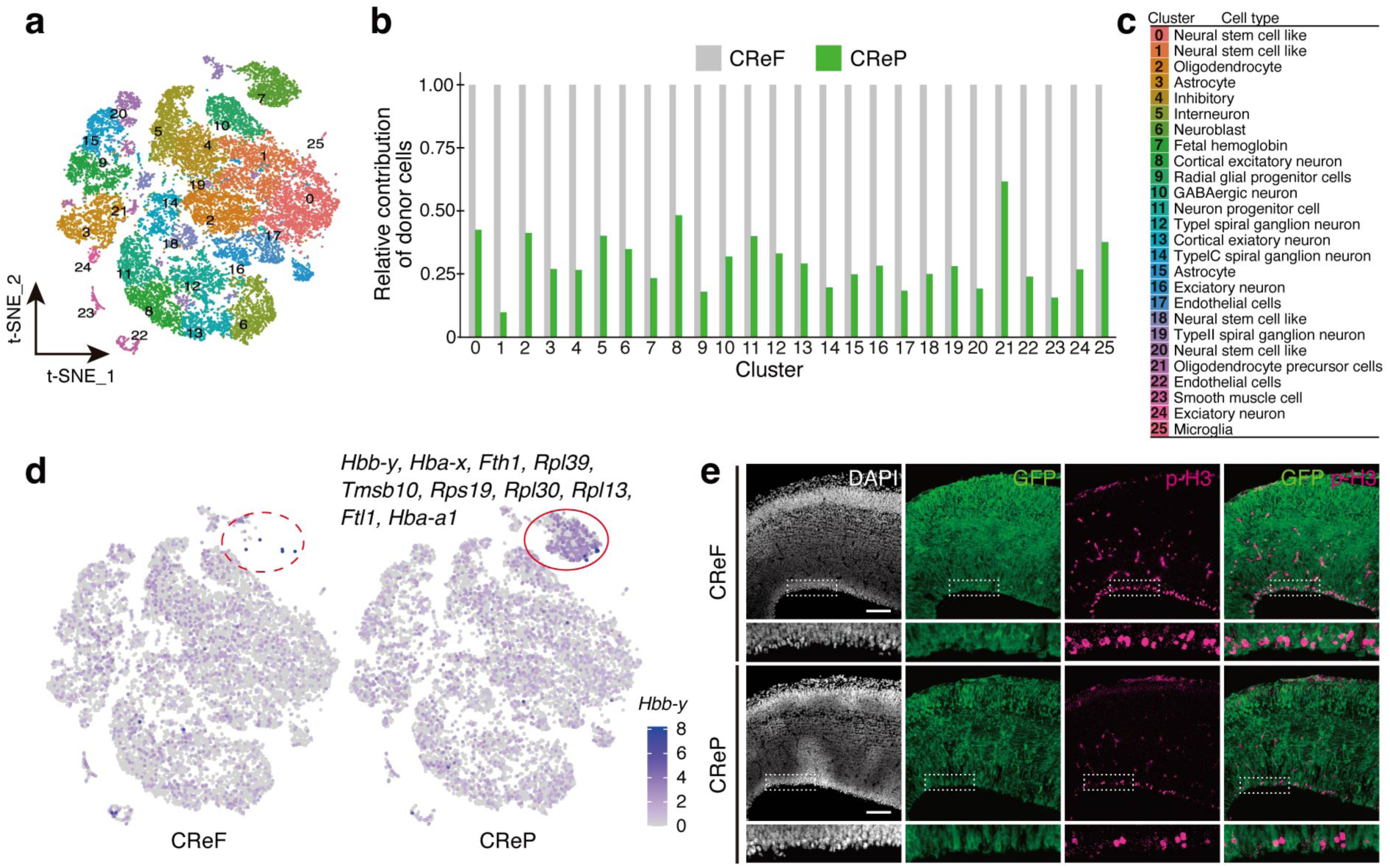
Distinct Cellular Profiles and Integration Deficits in CReP Brains Revealed by Single-Nucleus RNA Sequencing. (**a**) Cell clusters of CReP brains profiled by single nuclear RNA-sequencing (snRNA-seq) analysis. (**b**) Relative contribution rate of GFP-positive donor-derived cells per cluster; the contribution of fertilized embryo-derived cells in CReF is normalized to 1. (**c**) Cell types of each identified clusters. (**d**) Fetal hemoglobin population specifically identified in CReP brains (red circle). (**e**) Representative immunofluorescence images of CReP and CReF brains. Scale bar, 100 μm.

We also identified a CReP-specific population expressing embryonic globin genes (e.g. *Hbb-y and Hba-x*) (Fig. 3d). Given that embryonic globin expression is normally restricted to early embryonic stages before E11.5, this population likely reflects developmental delays in the CReP brain. Alternatively, this population may reflect inclusion of circulating erythroid or other vascular-associated cells in the brain preparation, for example due to blood–brain barrier immaturity or vascular leakage in CReP brains.

To validate at the tissue level the snRNA-seq–based inference of reduced neural stem/progenitor proliferation and maintenance, we performed histological analyses. This confirmed a marked reduction of phospho-Histone H3 (p-H3)-positive neural stem cells (NSCs) in the ventricular zone (VZ), accompanied by a significant loss of neural progenitors in the subventricular zone (SVZ) (Fig. 3e). Together with the marked decrease in Pg-derived contribution, these findings suggest cell-intrinsic defects in Pg-derived cells that compromise neural development and tissue integrity.

### Reduced proliferation and increased apoptosis in CReP-derived brain cells

To elucidate the mechanisms underlying the abnormalities in the CReP mouse brain, we cultured embryonic brain cells from E14.5 CReP and CReF embryos *in vitro*[28]. Both cultures expand over time; however, the fraction of GFP-positive donor-derived cells from fertilized embryos remained stable in CReF cultures, whereas the fraction of Pg-derived donor cells declined progressively in CReP culture after day 5 (Figs. 4a and 4b). Despite replicate-to-replicate variation in host to donor ratios, Pg-derived cells consistently showed reduced representation over time.

**Fig. 4.**
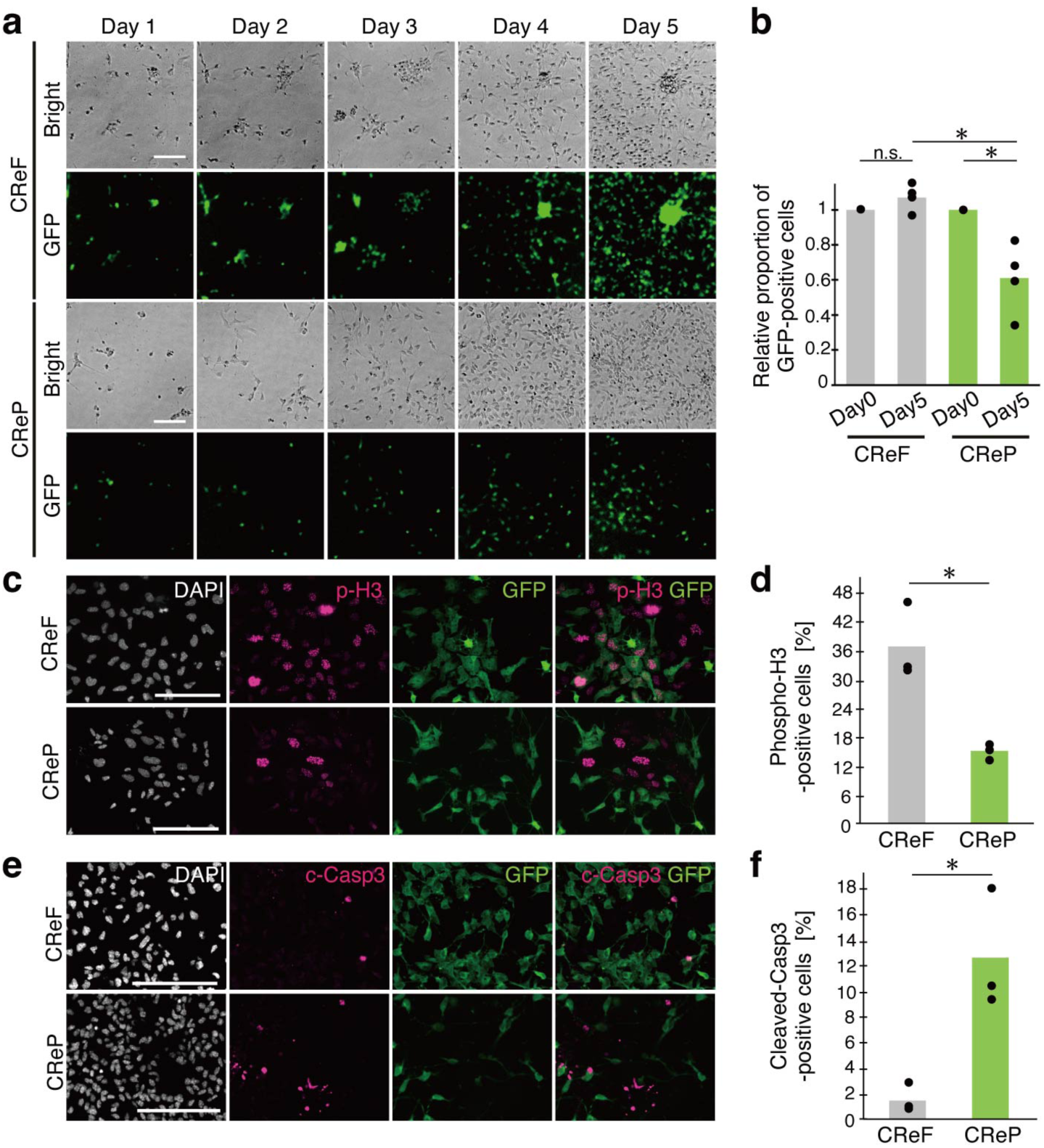
Reduced proliferation and increased apoptosis in Pg-derived brain cells *in vitro*. (**a**) Representative images of brain cells from CReP and CReF cultured from Day 1 to Day 5. Scale bar, 100 μm. (**b**) Relative GFP-positive proportion after five days of culture. The proportion at Day 0 was normalized to 1. Each dot represents an individual replicate. \**P*<0.05. (**c**) Representative immunofluorescence images of brain cells cultured for three days. Scale bar, 100 μm. (**d**) Quantification of p-H3-positive cells, representing proliferating cells. Each dot represents an individual replicate. \**P* < 0.05. (**e**) Representative immunofluorescence images of brain cells cultured for five days. Scale bar, 100 μm. (**f**) Quantification of cleaved caspase-3-positive apoptotic cells. Each dot represents an individual replicate. \**P*<0.05

We next quantified proliferative activity by p-H3 immunostaining and observed a marked reduction of mitotic cells in CReP cultures (Figs. 4c and 4d). Additionally, apoptosis was significantly elevated in CReP cultures, evidenced by an increased number of cleaved-Caspase3-positive cells compared to CReF cultures (Figs. 4e and 4f). These findings indicate that CReP-derived cells possess intrinsic proliferative deficiencies and heightened susceptibility to apoptosis which, together with in vivo snRNA-seq and histological analyses, provide a mechanistic basis for the diminished contribution of Pg-derived cells during midgestational brain development.

### Notch hyperactivation leads to skewed gliogenic differentiation in CReP brains

Prompted by the observed reduction in neural stem/progenitor proliferation in CReP brains, we examined signaling pathways regulating these processes, focusing on Notch. Notch signaling governs cell-fate decisions; in the embryonic brain, it sustains NSC self-renewal while constraining neuronal differentiation, and promotes gliogenesis (Fig. 5a)[29–33]. The paternally expressed imprinted gene, *Dlk1* modulates interactions between *Notch1* and its ligands[34,35]. RNA-seq analysis of CReP brains revealed reduced *Dlk1* expression in Pg-derived cells compared with fertilized embryo-derived cells in CReF brains, accompanied by increased expression of *Notch1*, its ligand *Dll1*, and the downstream effector *Hes1* (Fig. 5b). To assess Notch signaling activity, brain cells from E14.5 CReP and CReF embryos were cultured for five days and stained for Notch intracellular domains (NICD), which is generated upon ligand-dependent cleavage, translocate to the nucleus, and mediates downstream transcription. CReP-derived cells exhibited markedly stronger NICD signals, with frequent nuclear puncta, consistent with excessive Notch activation (Figs. 5c and 5d).

**Fig. 5.**
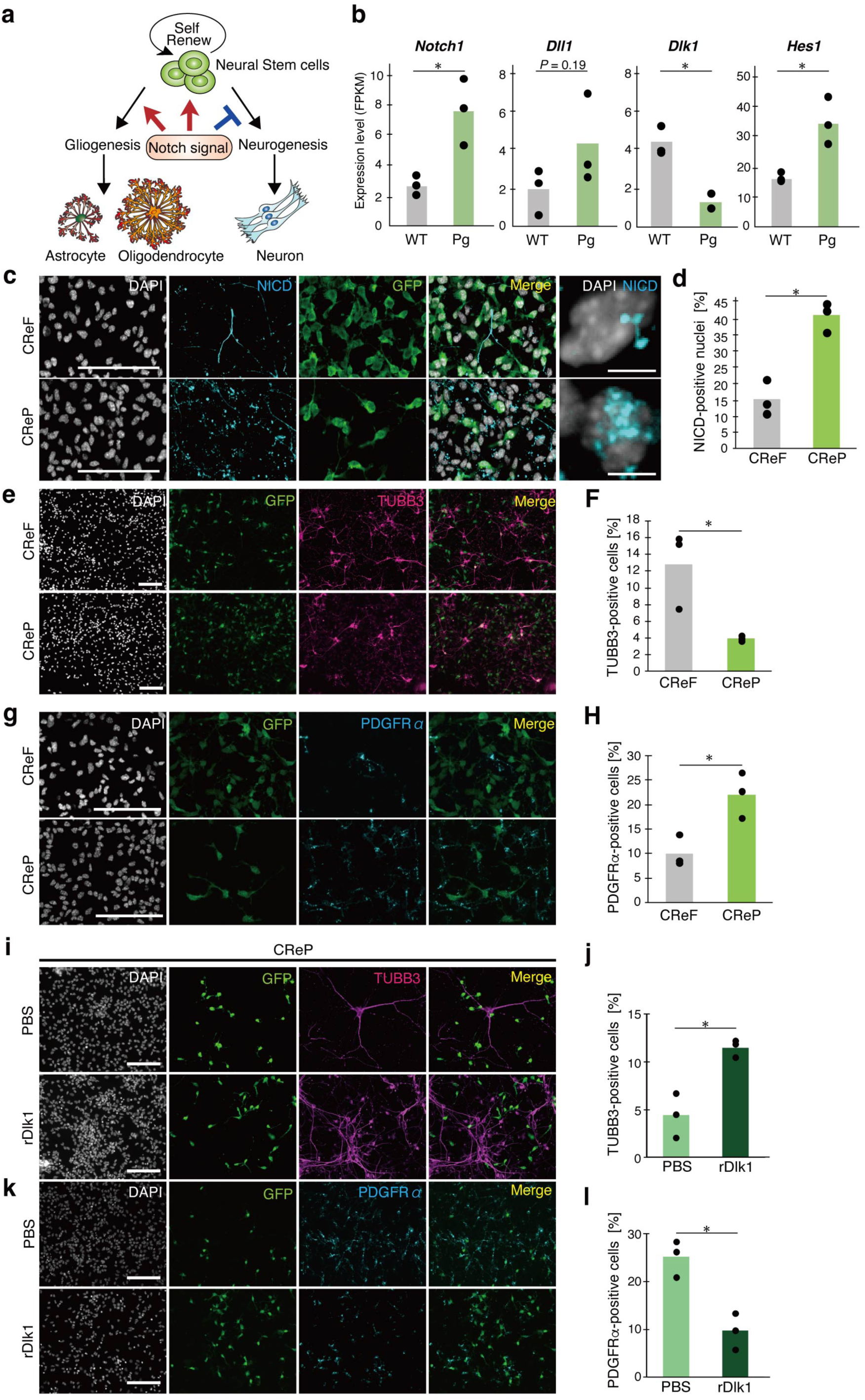
Notch hyperactivation with reduced Dlk1 in Pg-derived cells CReP brains skews fate toward gliogenesis. (**a**) Schematic of the Notch signaling pathway in neural stem cell fate determination. (**b**) Expression level of *Notch1, Dll1, Dlk1* and *Hes1* in GFP-positive sorted cells from CReF and CReP brains, as analyzed by RNA-sequencing. WILD-TYPE refers to fertilized embryo-derived cells from CReF embryos, and Pg refers to Pg embryo-derived cells in CReP embryos. Each dot represents an individual replicate. \**P*<0.05. (**c**) Representative immunofluorescence images of Notch intracellular domain (NICD) of brain cells cultured for five days. Scale bar, 100 μm. (**d**) Quantification of NICD-positive nuclei as a proportion of DAPI-positive cells in CReP and CReF cultures. Each dot represents an individual replicate. \**P*<0.05. (**e**) Representative immunofluorescence images of Tubulin beta III (TUBB3)-positive neurons in the brain cells cultured for five days. Scale bar, 100 μm. (**f**) Quantification of TUBB3-positive neurons as a proportion of DAPI-positive cells of CReP and CReF. Each dot represents an individual replicate. \**P*<0.05. (**g**) Representative immunofluorescence images of PDGFRα-positive oligodendrocyte precursor cells (OPCs) in the brain cells cultured for five days. Scale bar, 100 μm. (**h**) Quantification of PDGFRα-positive OPCs as a proportion of DAPI-positive cells of CReP and CReF. Each dot represents an individual replicate. \**P*<0.05. (**i**) Representative immunofluorescence images of TUBB3 in cultured CReP brain cells after 72 h of recombinant Dlk1 (rDlk1, 200 ng/mL) or PBS control treatment. Scale bar, 100 μm. (**j**) Quantification of TUBB3-positive neurons as a proportion of DAPI-positive cells in each condition. Each dot represents an individual replicate. \**P*<0.05. (**k**) Representative immunofluorescence images of PDGFRα in cultured CReP brain cells after 72 h of rDlk1 (200 ng/mL) or PBS control treatment. Scale bar, 100 μm. (**l**) Quantification of PDGFRα-positive OPCs as a proportion of DAPI-positive cells in each condition. Each dot represents an individual replicate. \**P*<0.05.

We next evaluated cell-fate specification *in vitro* by immunostaining after five days in culture of E14.5 CReP and CReF brains for Tubulin beta III (TUBB3; neurons), and Platelet-Derived Growth Factor Receptor alpha (PDGFRα; oligodendrocyte precursor cells, OPCs). The proportion of TUBB3-positive neurons was significantly reduced (Figs. 5e and 5f), whereas PDGFRα-positive cells increased (Figs. 5g and 5h) in CReP relative to CReF, indicating skewed differentiation toward gliogenesis with reduced neurogenesis.

To test the hypothesis that loss of *Dlk1* in Pg-derived cells drives Notch hyperactivation and gliogenic bias, we supplemented cultures with recombinant Dlk1 (rDlk1). rDlk1 increased the proportion of TUBB3-positive neurons relative to PBS-treated controls (Figs. 5i and 5j) and concomitantly reduced PDGFRα-positive OPCs (Figs. 5k and 5l). Together, these results demonstrate that loss of paternal Dlk1 expression leads to Notch signaling hyperactivation, thereby driving gliogenic bias and impaired neurogenesis in CReP brains.

## Discussion

This study established the CReP method as a generalizable platform to overcome the long-standing barrier of early post-implantation lethality in Pg embryos and to interrogate imprinting functions during later development. By creating tissue-specific niche in recipient embryos (e.g., *Wnt1* knockout for the brain) and complementing it with donor cells, CReP enables Pg-derived cells to survive and integrate into late-stage mouse embryos while rescuing organ-level morphological defects. Although chimeras composed of Pg and wild-type embryos have been reported, Pg-derived cell contribution has varied unpredictably across embryos, limiting their utility for research[36]. In contrast, CReP enables targeted Pg-cell contribution to specific tissues simply by altering the knockout locus in the recipient, facilitating the generation of CReP mice across organs and stages. Systematic datasets generated across diverse CReP contexts will substantially advance studies of genomic imprinting.

Despite regional rescue via mosaic integration of fertilized embryo-derived recipient cells, CReP embryos remained growth-restricted and most neonates did not survive, indicating organism-wide effects of imprinting defects beyond the complemented tissue. Notably, severe phenotypes persisted despite mosaic recipient-cell integration, paralleling imprinting disorders, such as Prader-Willi syndrome, Kagami–Ogata syndrome, and Silver–Russell syndrome, which can feature mosaic uniparental disomy alongside normal karyotype cells[37–39]. Our findings suggest that imprinting abnormalities shape tissue development through both cell-autonomous and non-cell-autonomous influences, exemplified here by signaling alteration in the Notch pathway. Furthermore, variation in mosaicism rates among CReP mice may help recapitulate the clinical heterogeneity of mosaic imprinting disorders.

CReP also offers an orthogonal route to identify previously uncharacterized imprinted genes at late developmental stages. Conventional discovery has relied on allele-biased expression using single-nucleotide polymorphisms (SNPs) in hybrid embryos, a strategy inapplicable to loci lacking informative SNPs[40]. In contrast, DEGs in Pg-derived cells within CReP brains can flag candidates independent of SNPs. Approximately 20% of previously reported imprinted genes were recovered as DEGs in our analysis, supporting this concept, and the candidate list likely includes late-stage, brain-specific regulators. Follow-up with allele-specific expression, reciprocal crosses, and locus-resolved epigenomic profiling will be required to assign imprinting status definitively.

At the mechanistic level, our data implicate the Dlk1–Notch axis in Pg-cell dysfunction in CReP mice. In the embryonic brain, Notch signaling regulates NSC self-renewal and restricts neuronal differentiation[29]. The paternally expressed, non-canonical Notch ligand Dlk1 modulates interactions between Notch1 and its ligands[41]. In CReP brains, showed reduced *Dlk1* expression together with increased NICD staining and elevated *Notch1/Dll1/Hes1*, consistent with Notch hyperactivation[42–45]. Correspondingly, NSCs and progenitor pools were diminished in vivo, and in vitro Pg-derived cells exhibited fewer mitoses and more apoptosis. Fate-marker analyses further indicated skewed differentiation toward gliogenesis with reduced neurogenesis; recombinant Dlk1 partially increased neuronal output and reduced OPC frequency, supporting a causal link between reduced Dlk1 and Notch hyperactivation. Prior zebrafish work showing the requirement for Notch down-regulation during OPC differentiation aligns with these observations[46,47] . Together, the data highlight a paternal-allele–encoded regulatory role for Dlk1 in calibrating Notch activity to maintain the timing and balance of neural cell production (Fig. 6). Given reported association between reduced NSC numbers and neurodegenerative disorders such as Alzheimer’s disease[48] or neuropsychiatric conditions like schizophrenia[49], longitudinal studies of adult CReP mice may yield critical insights into the pathogenesis of these conditions.

**Fig. 6.**
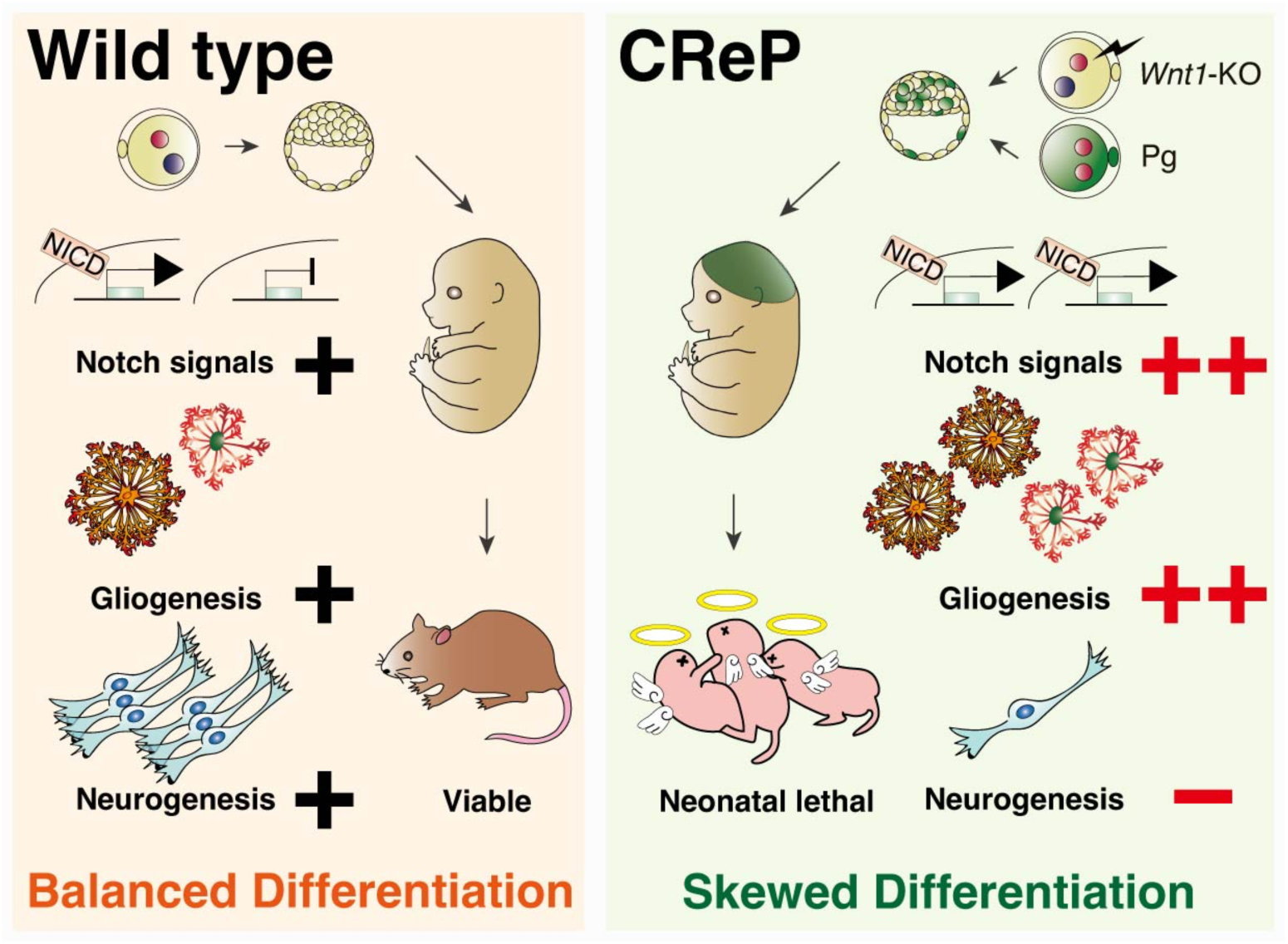
Schematic model: Dlk1 dosage constrains Notch signaling to maintain neural cell fate balance in the developing mouse brain.

In conclusion, our CReP model moves imprinting research beyond the implantation bottleneck. Through simple modification of recipient knockouts, it enables tissue-targeted analyses and nominates candidate imprinted regulators without reliance on SNPs. By combining organ-targeted complementation with cell-state–resolved readouts and targeted pathway perturbations, this conceptual framework offers a clear approach to elucidate how parent-of-origin information orchestrates tissue growth, lineage allocation, and organismal viability during later development.

Schematic model. In WILD-TYPE contexts, paternally expressed Dlk1 restrains Notch signaling to balance neural cell-type specification. In CReP, reduced Dlk1 causes Notch hyperactivation, leading to skewed cell-fate specification toward gliogenesis accompanied by reduced neurogenesis, growth restriction, and neonatal death shortly after birth.

## Material and Methods

### Animals

All animal experiments were performed in accordance with the guidelines of the Animal Care and Use Committee of the Graduate School of Frontier Biosciences, Osaka University and the Animal Care and Use Committee of Toho University (ethical approval protocol 22-51-518). F1 mice from C57BL/6NCrSlc × DBA/1JJmsSlc crossings (B6D2F1), ICR, and vasectomized ICR mice were purchased from Sankyo Labo Service Corporation (Tokyo, Japan). F1 hybrid mice generated by crossing Rosa26 Cre-reporter knock-in mice (R26GRR mice)[50] with DBA/1JJmsSlc mice were used for Pg embryo in this study. The R26GRR mouse line was originally developed and distributed by Ken-ichi Yagami (University of Tsukuba, Tsukuba, Japan) and RIKEN BRC (Tsukuba, Japan). The R26GRR mice used for breeding were kindly provided by Prof. Hiroshi Sasaki (Osaka University, Osaka, Japan) and Dr. Masakazu Hashimoto (Fukushima Medical University, Fukushima, Japan). All mice were housed under 12-hour light/12-hour dark cycle with free access to food and water.

### Superovulation and Embryo Collection

Superovulation was induced in female B6D2F1 (≥21 days) by intraperitoneal injection of CARD Hyper Ova (Kudo), followed 48 h later by 7 U of hCG (Aska Pharmaceutical).

Females were then mated with male B6D2F1 (≥15 weeks). For Pg activation, the females were not mated. Mice were euthanized 18 h after hCG injection, and fertilized embryos were recovered from the ampulla of oviduct into M2 medium (Sigma) supplemented with 4 mg/mL BSA (EMD Millipore) (M2+BSA) and 1 mg/mL hyaluronidase (Sigma). Embryos were washed in M2+BSA and cultured in KSOM (ARK Resource Co., Ltd.) until use.

### CRISPR/Cas9-Mediated Gene Editing

Single guide RNAs (sgRNAs) were synthesized by *in vitro* transcription (IVT). DNA templates were generated by PCR from px330 plasmid (Addgene) using primers incorporating the T7 promoter and *Wnt1* target sequences (Supplementary Table S1). IVT was performed using the MEGAshortscript T7 Kit (Invitrogen). The electroporation (Ep) solution comprised three *Wnt1*-sgRNAs (each at 100 ng/μl) in Opti-MEM (Gibco) and 200 ng of Cas9 protein (Thermo Fisher Scientific). Fertilized embryos were rinsed in Opti-MEM and Ep solution, aligned between electrodes in 5 μl Ep solution, and electroporated (25 V, 3 ms ON/ 97 ms OFF, for 5 cycles). Embryos were immediately washed in M2+BSA and cultured in KSOM.

### Parthenogenetic Activation and Chimera Generation

Unfertilized oocytes were collected from superovulated Rosa-B6D2F1 mice as above, incubated in KSOM for 1 h, then in KSOM containing 2 mM EGTA (Bio Medical Science), 5 mM SrCl2, and 5 μg/mL cytochalasin B for 5–6 h at 37 °C[51]. After confirmation of pronuclear formation, embryos were washed and further cultured in KSOM. Aggregation wells were prepared in 6 cm dishes using aggregation needles (BLS); each well was filled KSOM drops, overlaid with mineral oil, and equilibrated overnight in a CO₂ incubator. Zona pellucida were removed from fertilized and parthenogenetic embryos at ∼42 h post-electroporation or parthenogenetic activation using 5 mg/mL PRONASE (Millipore). One fertilized and one parthenogenetic embryo were placed in a single aggregation well and co-cultured for 48 h at 37°C. Pseudopregnant ICR females (>6 weeks) were obtained by mating with a vasectomized ICR males; vaginal plugs were checked the next day (defined as day 0). Under mixed anesthesia, chimera blastocysts were transferred into the uterine on day 2.5 using a glass pipette.

### Fetal Brain Cell Collection

Dissection and culture of fetal brains were performed as previously described[28]. For bulk RNA sequencing, GFP-positive cells were sorted using a FACS Aria III Cell Sorter (BD Biosciences) into Trizol LS reagent (Ambion, 10296-010). For single-nucleus RNA sequencing, cells were preserved in Cell Reservoir One with DMSO (Nacalai tesque). Three brain cell preparations with similar GFP-positive proportions were pooled into one sample. For longitudinal culture measurements, cells were detached using Accutase (Sigma) and the GFP-positive fraction was quantified on a CytoFLEX flow cytometer (Beckman Coulter) from seeding to day 5. Cells cultured on coverslips were fixed with 4% paraformaldehyde (PFA) for 10 minutes at room temperature on day 3 and 5. For Dlk1 supplementation, brain cells were treated for 72 h with recombinant Dlk1 (R&D systems, 1144-PR-025/CF, 200ng/ml) or PBS, then fixed in 4% PFA for 10 min.

### RNA Sequencing Analysis

Total RNA from sorted cells in Trizol LS was purified using the Direct-zol RNA MicroPrep kit (ZYMO Research). Library was sequenced (101-bp single-end) on an Illumina HiSeq 2500. Reads were mapped to the mouse reference genome (mm10) using TopHat (v2.0.13), with Bowtie2 (v2.2.3) for alignment and SAMtools (v0.1.19) for processing. FPKM values were calculated using Cufflinks (v2.2.1). For differential expression analysis, RNA-seq count data were processed using the edgeR package in R. Normalization was performed using the TMM method, and log-transformed counts per million (CPM) were calculated. p-values were adjusted using the Benjamini-Hochberg (BH) method, and the results were exported for further analysis. GO enrichment analysis was performed using the clusterProfiler package in R.

### Single-Nucleus RNA Sequencing

Single-nucleus RNA sequencing was performed by Rhelixa Inc. using 10x Genomics platform for libraries preparation. Sequencing on a NovaSeq 6000 yielded ∼400 million pair-end reads.sData processing involved applying the following filtering thresholds: nFeature_RNA > 500, nCount_RNA > 500, nCount_RNA < 50,000, and percent.mito < 20%.

### Immunofluorescence

Brains were fixed in 4% PFA for 3 h at room temperature, washed three times in PBS, transferred to 30% sucrose/PBS until sinking, embedded in O.C.T. compound (Sakura Finetek Japan Co., Ltd.) for cryopreservation. Cryosections were prepared at a thickness of 10 µm. For coverslip cultures, fixed cells were blocked with Blocking One (Nacalai tesque), permeabilization in 0.4% phosphate buffered saline with Tween 20 (PBST) for 30 min at room temperature, washed five times in PBS, cells were incubated with primary and secondary antibodies listed in Supplementary Table S5. Nuclei were counterstained with DAPI (Sigma, D9542, 1 mg/ml) and samples were mounted with antifade reagent (Nacalai tesque). Images were acquired on a Keyence, BZ-X710). At least 1000 cells were counted for each condition. Two-sided t-tests were performed for each dataset with a significance level of α = 0.05. For cryosection, sections were blocked in 10% donkey serum in 0.3% PBST for 1h at room temperature, incubated with primary antibodies overnight at 4℃, followed by secondary antibodies for 1h at room temperature. We used primary and secondary antibodies listed in Supplementary Table S5. Confocal images were acquired on an Olympus FV3000 CellSens.

## Abbreviations

Pg: parthenogenetic
Ag: Androgenetic
CReP: Cell replacement with parthenogenote-derived cells
CReF: Cell Replacement with Fertilized embryo-derived cells
DMRs: DNA methylation at differentially methylated regions
KO: Knockout
FACS: Fluorescence-activated cell sorter
MEGs: Maternally expressed genes
PEGs: Paternally expressed genes
Non-IGs: Non-imprinted genes
DEGs: Differentially expressed genes
NSCs: Neural stem cells
VZ: Ventricular zone
SVZ: Subventricular zone
NICD: Notch intracellular domains
TUBB3: Tubulin beta III
PDGFRα: Platelet-Derived Growth Factor Receptor alpha
OPCs: Oligodendrocyte precursor cells
SNPs: Single-nucleotide polymorphisms

## Data availability

All sequencing data (bulk and single-nucleus RNA sequencing) generated in this study have been deposited in the NCBI BioProject database under accession number PRJNA1334549.

## Ethics statement

The animal study was reviewed and approved by the Animal Care and Committee of the Graduate School of Frontier Biosciences, Osaka University and the Toho University.

## Author contribution

MT and SY conceived, designed, and coordinated the study. MT mainly performed experiments and acquired the data. YZ, ZL and LS analyzed snRNA-sequencing data. YZ and KM stained pH3-positive cells in cryosection. SY and TN supervised. MT and SY wrote the manuscript. All authors contributed to the interpretations and conclusion of the study.

## Acknowledgements

We acknowledge the NGS Core Facility of the Genome Information Research Center at the Research Institute for Microbial Diseases, Osaka University, for their support in RNA sequencing. We also thank Ken-ichi Yagami (University of Tsukuba) and RIKEN BRC through the National Bio-Resource Project of the MEXT, Japan, for providing R26GRR mice.

## Declaration of generative AI and AI-assisted technologies in the writing process

During the preparation of this work, the authors used OpenAI ChatGPT solely for English language editing (spelling, grammar, and minor clarity). After using this tool, the authors reviewed and edited the content as needed and take full responsibility for the content of the published article.

