## Supplementary Figures for "Parthenogenote-Derived Brain Unveils the Critical Role of Paternal Genome in Neural Development"

**Supplementary Fig. S1-4**

**Supplementary Table S1-5**

**Supplementary Fig. S1**

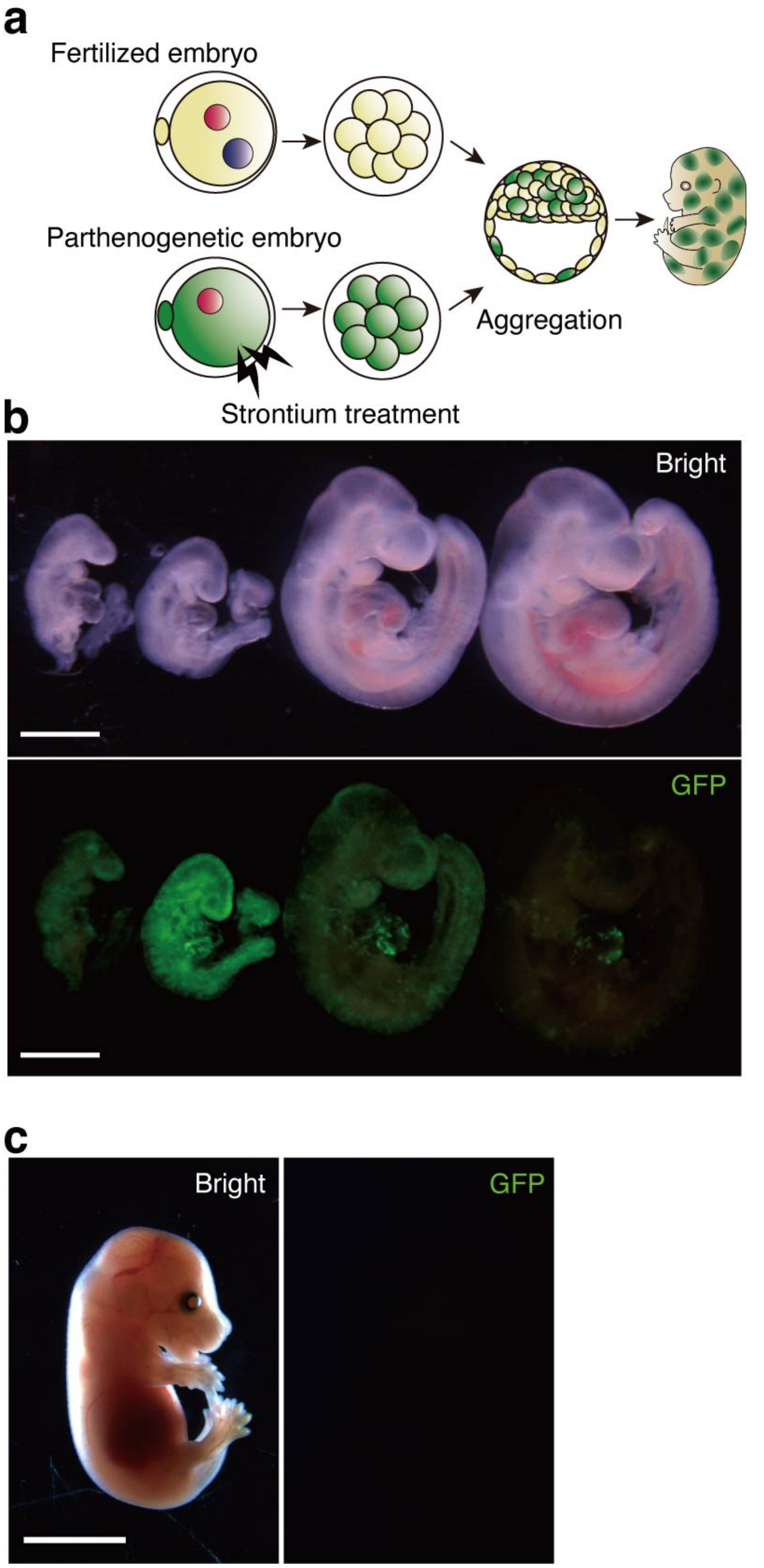

**Supplementary Fig. S1 Chimeras of fertilized and parthenogenetic embryos**

(**a**) Experimental scheme for generating chimeras of fertilized and parthenogenetic (Pg) embryos. (**b**) Representative E9.5 chimeric embryos; GFP-positive cells indicate the contribution of Pg embryos. Scale bars, 1mm. (**c**) Representative E14.5 chimeric embryos derived from fertilized and Pg embryos. Scale bars, 1mm.

**Supplementary Fig. S2**

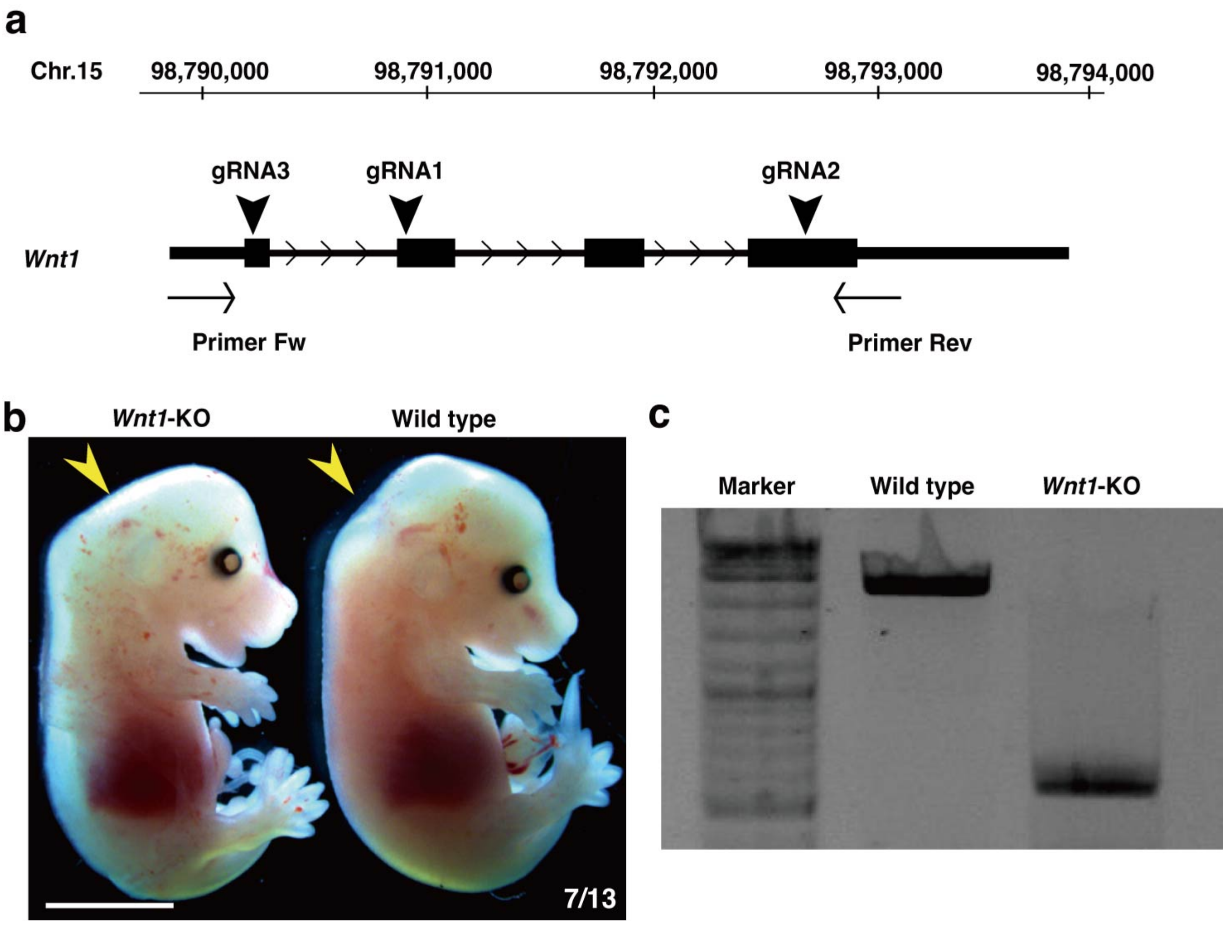

**Supplementary Fig. S2 CRISPR-Cas9-mediated *Wnt1* knockout**

(**a**) Schematic showing guide RNA target sites within the *Wnt1* locus and primers used for genotyping. (**b**) Representative E14.5 *Wnt1*-KO and wild-type embryos. Yellow arrowheads indicate the characteristic forebrain/midbrain defects in *Wnt1*-KO embryos. Scale bar 4 mm. (**c**) Genotyping PCR using Primer-Fw and Primer-Rev confirms a large deletion in *Wnt1*-KO embryos.

**Supplementary Fig. S3**

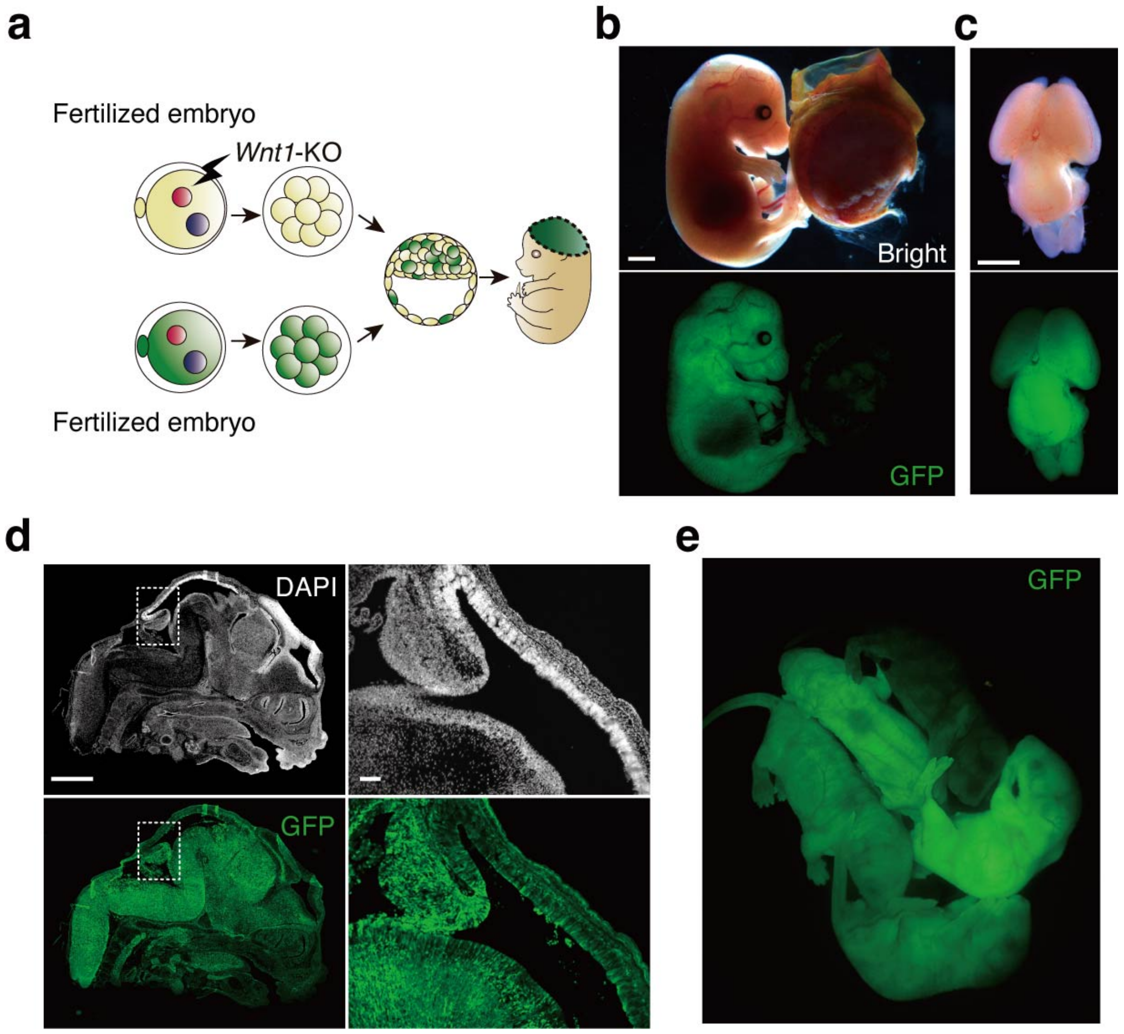

**Supplementary Fig. S3 Development of CReF mice**

(**a**) Experimental scheme for establishing Cell replacement with fertilized (CReF) embryos. (**b and c**) Representative E14.5 CReF embryos (**b**) and their brains (**c**). GFP indicates contribution from fertilized embryos. Scale bars, 1mm. (**d**) Representative cryosections from E14.5 CReF embryo. The enlarged section is indicated by the dotted rectangle. Scale bar 1mm. (**e**) Representative CReF neonates at postnatal day 0.

**Supplementary Fig. S4**

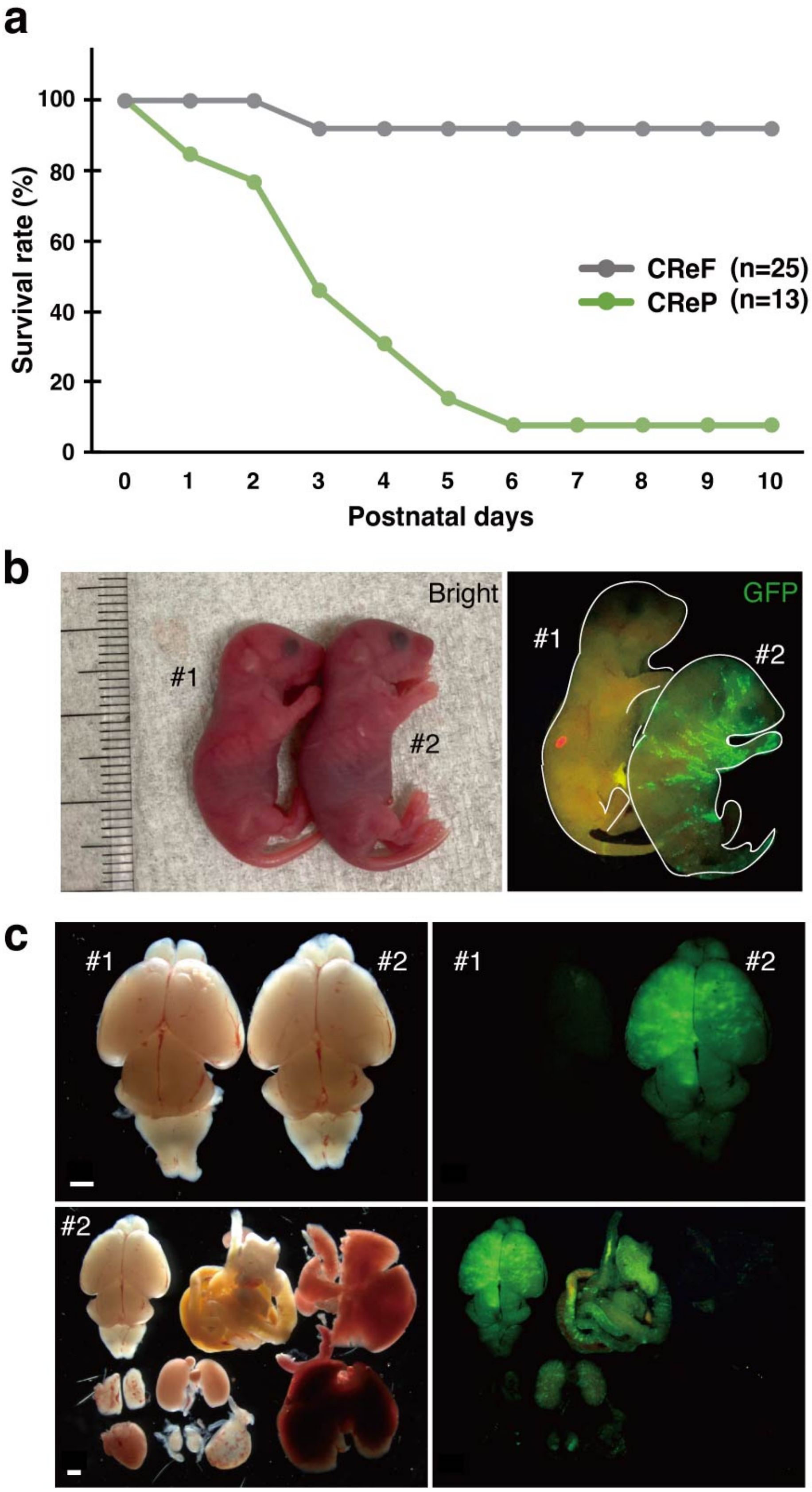

**Supplementary Fig. S4 Neonatal lethality of CReP mice**

(**a**) Survival rates of CReP and CReF mice. (**b**) Representative images of CReP mice at postnatal day 0. Contribution of GFP-positive Pg cells is detectable in mouse #2. (**c**) Representative images of CReP mice organs at postnatal day 0.

**Supplementary Table S1**

Sequences of sgRNAs used for *Wnt1* knockout by CRISPR-Cas9.

**Supplementary Table S2**

Differentially expressed genes in CReP.

**Supplementary Table S3**

Results of gene ontology analysis.

**Supplementary Table S4**

Cluster marker genes.

**Supplementary Table S5**

Antibodies used in this study.
